# Integrating Network Pharmacology And Experimental Verification To Explore The Mechanism Of Qionggui Power Against Atherosclerosis

**DOI:** 10.1101/2022.11.01.514795

**Authors:** Yuqing Wang, Yonghong Man, Xu Jiang, Dong Shui, Qing Zhao, Shujiao Li, Guo Zhuang

## Abstract

Qionggui Power (QP), a classic prescription in Traditional Chinese Medicine (TCM), has shown potential in the treatment of atherosclerosis during the past decades. However, the mechanism that mediates these cardiovascular benefits remains to be fully elucidated. Here, we investigated the effects and mechanisms of QP against atherosclerosis with network pharmacology approaches and *in vitro* model. The active ingredients and related targets of QP were collected from public databases. The hub targets and signaling pathways of QP against AS were defined by extensive application of bioinformatics approaches, including the protein-protein interaction (PPI) network, Gene Ontology (GO), and Kyoto Encyclopedia of Genes and Genomes (KEGG). The predicted major targets were validated in LPS-stimulated murine macrophages RAW264.7. The anti-inflammatory properties of QP were also evaluated in this model. *In silico* investigation of QP resulted in the identification of 18 active ingredients and 49 chemical targets intersecting with AS-related genes. And KEGG pathway analysis revealed a high enrichment in the Lipid and Atherosclerosis pathway of these chemical targets. Biochemical analysis showed marked effects of QP on the expression of predicted chemical targets (PPARr, CAT, PTGS2) and LPS-induced inflammatory genes (IL1, IL6, and TNFα). And these inhibitory effects were linked to the suppression of the NF-κB signaling pathway, which was activated by the LPS stimulus. Our findings revealed the therapeutic potential of QP in the prevention and treatment of atherosclerosis.

**Graphical Abstract:** 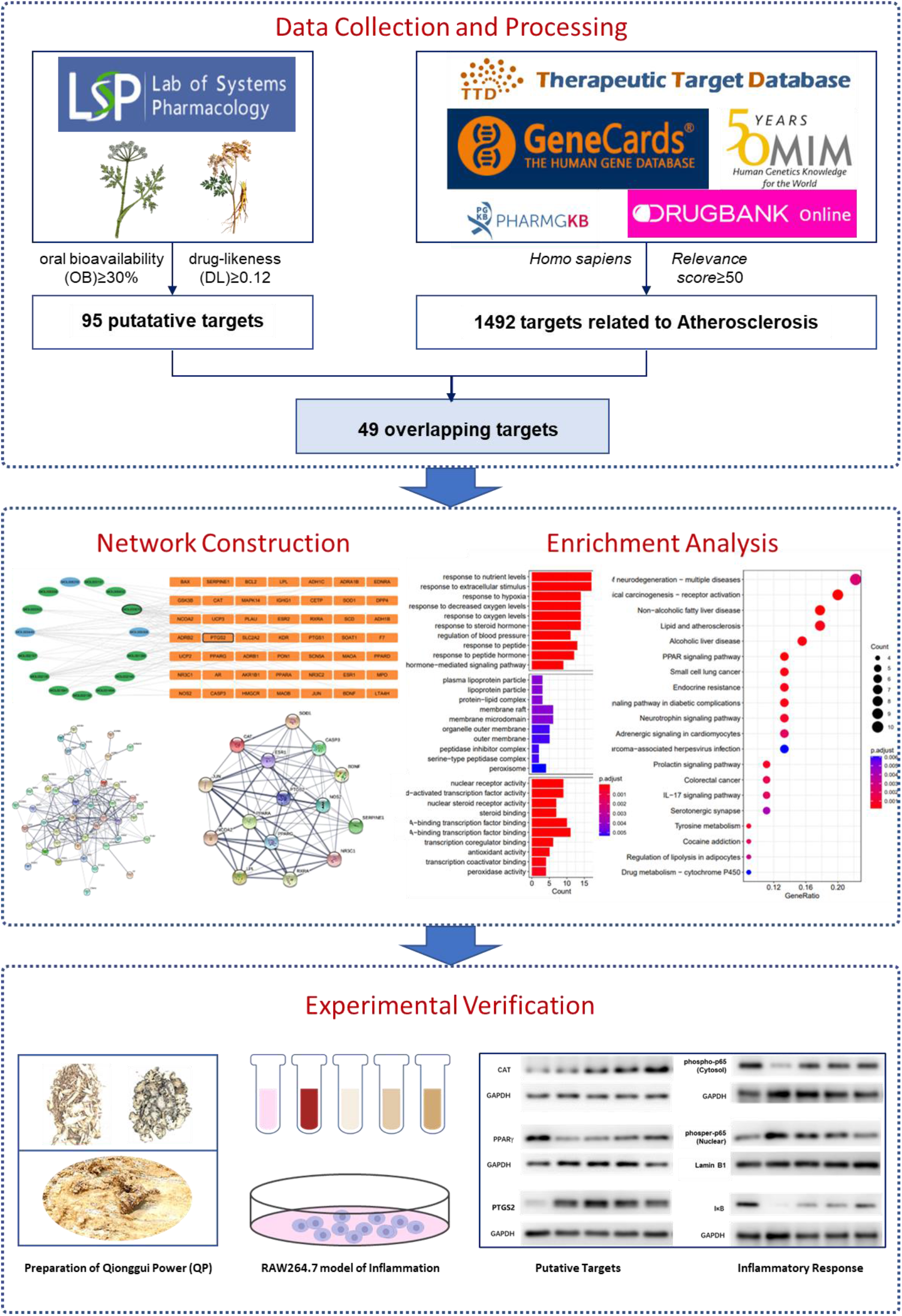

## 1. Introduction

cardiovascular disease (CVD) is the leading cause of mortality ^[1, 2]^. Globally, CVD causes an estimated 17 million deaths, accounting for one-third of all deaths annually ^[3]^. Risks of developing CVD, including aging, family history, high blood pressure, high cholesterol, obesity, diabetes, and smoking ^[4]^, are tightly related to the altered structure of arterial intima ^[5]^ and the modification of plasma-derived lipoprotein ^[6]^. These alterations accumulate plaque or even form a thrombus in the artery wall, which induce a complex process called atherosclerosis (AS) ^[7]^. Among the catalog of molecular mechanisms underlying AS, lipid-driven inflammation establishes a central role in the pathogenesis of AS ^[8]^. Intensified inflammatory activation may lead to local proteolysis, plaque rupture, and thrombus formation, wherein the pro-inflammatory mediates secreted by recruiting macrophages, such as IL1, IL6, and TNFα, contribute to the local inflammation and growth of the plaque ^[8]^. Therefore, inflammatory markers are devoted to the identification of active chemical components and medical potential for AS treatment.

Qionggui Power (QP) is composed of two herbs, Chuanqiong (*Ligusticum chuanxiong hort*) and Danggui (*Angelica sinensis*). The dried rhizomes of these herbs have long shown clinical benefits in gynecological diseases. In recent years, several experimental studies have evaluated the cardioprotective effects of QP using modern scientific methods ^[9]^, and these investigations raise the possibility of its application in AS treatment. Although modern pharmacological experiments have suggested various activities associated with the anti-inflammatory, anti-oxidative, and anti-aging effects of QP ^[10, 11]^, the bioactive component and pharmacological mechanism underlying these benefits remain obscure.

Network pharmacology establishes the foundation of a new paradigm in drug discovery. This method shows significant implications in tackling the two major sources of attrition in drug development - efficacy and toxicity ^[12]^. Therefore, network pharmacology provides an effective approach to validate both target combinations and optimize multiple structure-activity relationships while maintaining drug-like properties ^[13]^. Using this paradigm, the holistic philosophy of Traditional Chinese Medicine (TCM) is contextualized by pharmacologic mechanisms, wherein an interacting network between multiple components, drug targets, and pathways is defined efficiently ^[14]^.

In this study, the ingredients and drug targets of QP were obtained from a public database, Traditional Chinese Medicine Systems Pharmacology (TCMSP). The screening criteria for active ingredients were set as oral bioavailability (OB)⩾30% and drug-likeness (DL)⩾0.12. A total of 49 molecular targets against AS of these chemicals were identified using the PPI network. These molecular targets underwent functional enrichment using GO and KEGG analyses, which revealed a strong involvement in lipid and atherosclerosis pathways. Furthermore, 15 hub targets were obtained by CytoNCA, a Cytoscape plugin integrating calculation, evaluation, and visualization analysis for multiple centrality measures, and the screening conditions included BC, CC, DC, EC, LAC, and NC. The major proteins involved in the predicted pathway and signaling pathways were validated in an LPS-induced inflammatory model with a macrophage cell line (RAW264.7) in vitro.

## 2. Materials and Methods

### 2.1. Screening for Active ingredients and related targets from QP

The active ingredients and related targets were screened from the pharmacology platform of Chinese herbal medicines, Traditional Chinese Medicine Systems Pharmacology (https://tcmsp-e.com/tcmsp.php, TCMSP) by using the keywords ‘Chuangqiong’ and ‘Danggui’. And the screening criteria for an active ingredient were set as oral bioavailability (OB)⩾30% and drug-likeness (DL)⩾0.12. All the related targets’ names were standardized based on the data from the UniProt database.

### 2.2. Establishing a collection of AS-related genes

A collection of genes that are linked to AS pathogenesis and therapeutics were established based on data from five public genetic and clinical databases, including GeneCards (https://www.genecards.org/), OMIM (https://omim.org/), PharmGKB (https://www.pharmgkb.org/), TTD (http://db.idrblab.net/ttd/), Drugbank (https://go.drugbank.com/). A total of 1492 AS-related genes were identified after removing the duplicate data.

### 2.3 Mapping for the targets of active ingredients and AS-related genes

Both the targets of active ingredients from QP and the AS-related genes were inputted into Cytoscape, an open-source software platform, to visualize the complex networks, and a mapping of these targets and active ingredients was constructed with integrated topological parameters. These parameters are tightly related to the importance of genes in AS pathogenesis or therapeutics.

### 2.4 Constructing Protein-Protein Interaction (PPI) Network and hub proteins network

The identified genes from the target mapping were imported into String database (https://string-db.org/) to build the Protein-Protein Interaction (PPI) Network. The minimum required interaction score for medium confidence is set as 0.4. After screening out the disconnected nodes in the network, short tabular text, which includes lists of one-edges, was derived and inputted into Cytoscrape. Then the hub target proteins were obtained with a Cytoscape plugin, CytoNCA, upon a combined screening of six topological factors: Betweenness centrality (BC), Closeness Degree (CD), Degree centrality (DC), Eigenvector centrality (EC), local average connectivity-based method (LAC), and network centrality (NC). And the screening criteria were set as greater than or equal to the median value for each of these factors.

### 2.4 Functional Enrichment Analyses

The name of potential target genes was transformed into the form of entrezID. Then the Kyoto Encyclopedia of Genes and Genomes (KEGG) and Gene Ontology (GO) enrichment analyses were performed with the R software for statistical computing and graphics. And the top 30 entities of KEGG analysis were mapped as bar plots, whereas the top 10 entities of GO analysis were mapped as bubble plots.

### 2.5 Preparation of Qionggui Power (QP)

Raw herbs were purchased from a local supplier, Henan Zhang Zhongjing Pharmacy Co. (Henan, China). The roots of *Ligusticum chuanxiong hort* were grown in Gansu province, China, and the roots of *Angelica sinensis* were grown in Xichuan province, China. Both herbs were dried and cut in the place of origin after harvest. The voucher specimens of Chuanqiong (2021-1117b) and Danggui (2021-1118b) were deposited in the museum of the Scientific Research Center, the Nanyang Medical College.

The aqueous extract of QP was prepared as follows. *Ligusticum chuanxiong hort* and *Angelica sinensis* were mixed in 2.33:1 weight ratio and macerated in 10-fold of water for 1.5 hours. The mixture was extracted at 100°C for 1h, and the residues were then subjected to two more subsequent extractions for 1 hour and 30 minutes, respectively. The water extract was concentrated by rotary evaporation and dried by vacuum freezing. The preparation of QP is shown in figure 4A. The extraction yield of the herbal formula QP was approximately 36%.

### 2.6 Cell culture

The mouse macrophage cell line RAW264.7 (ATCC number: TIB-71) was obtained from the cell bank of the Chinese Academy of Sciences (Beijing, China). Cells were cultured in a growth medium, consisting of Dulbecco’s modified Eagle’s medium (DMEM), with 17.5mM D-Glucose, 15.1 mM 4-(2-hydroxyethyl)-1-piperazine ethane sulfonic acid (HEPES), 2.5mM glutamine, 0.5 mM sodium pyruvate (Dalian Meilun Biotechnology Co., Dalian, China), supplemented with 10% v/v fetal bovine serum (FBS) (Tianhang Bio, Hangzhou, China) and 1% (v/v) penicillin/streptomycin sulfate cocktail at 37 °C in a humidified atmosphere containing 5% CO_2_. When cells reached full confluence, they were seeded in plates at a density of 4×10^5^/cm^2^. The ratio of medium volume to plate surface area is 0.3125ml/cm^2^, and the incubation medium consists of DMEM supplemented with 1% FBS. Then, the cells were allowed to reattach and settle for 12h in an incubator at 37°C. Subsequently, the cells were subjected to incubation for indicated periods under the following experimental conditions: (a) incubation medium alone; (b) 100ng/ml LPS (Beyotime Institute of Biotechnology, Haimen, China); (c) 100ng/ml LPS and 0.25mg/ml QP; (d) 100ng/ml LPS and 0.5mg/ml QP; (e) 100ng/ml LPS and 1mg/ml QP.

### 2.7 Formazan formation assay

The effects of QP on the viability of cells were assessed by formazan formation assay, wherein Cell Counting Kit-8 (Dalian Meilun Biotechnology Co., Dalian, China), a water-soluble tetrazolium salt (WST)-based assay was used. At the end of the indicated time for incubation, cells in a 96-cells plate were fed with an incubation medium containing 10μl/well WST-8 (10% Cell Counting Kit-8 solution). After incubation at 37 °C for 1h, Absorbance was read at 450 nm by a microplate spectrophotometer (Bio Tek, US).

### 2.8 RNA extraction, cDNA synthesis

At the end of the incubation period, total RNA was isolated from cultured cells in six-well plates using an Ultrapure RNA Kit (Jiangsu Kangwei century Biotechnology Co., Ltd, China) according to the manufacturer’s instructions. In brief, the culture medium was removed completely, and 1ml TRIZOL was added to each well. The plate was shaken with a circumferential shaker for 20min. Then, the lysis solution was collected and mixed with 200ul chloroform. After a violent vortex, the mixture was centrifuged at the speed of 12000g for 10 minutes at 4 °C. A total of 420ul aqueous-phase supernatant was collected and mixed with an equal volume of 70% ethanol. The mixture was then mounted into Spin Columns with a silica cartridge for purification. The purified RNA products were measured using a nanodrop spectrophotometer (Thermo Fisher Scientific, USA, 260 nm and A260/280 =1.8-2.0). A commercial cDNA synthesis kit, PrimeScript™ RT reagent Kit with gDNA Eraser (Takara Biomedical Technology Co., Ltd. Japan) was used for reverse transcription. The possible DNA contamination was cleared by DNase treatment. The cDNA synthesis temperatures were considered as follows: incubated at 37 °C for 15 min, 85 °C for 5 seconds, and finally kept at 4 °C.

### 2.9 Quantitative polymerase chain reaction (qPCR)

qPCR was performed with a commercial 2× Real-Time PCR master mix including SYBR Green I, LOW ROX (Code No. RR820, Takara, Japan). A standard PCR mixture contained 2.0ul template cDNA, 10μl TB Green Premix Ex Taq II, 0.8μl forward primer, 0.8μl reverse primer, 6.0μl RNase-free water, and 0.4ul ROX Reference Dye II. The cycling protocol included 95 °C (20 Sec), followed by 40 cycles of 95 °C (3 Sec) and 60 °C (30 Sec for annealing temperature). Additionally, a melt curve was used to check the specificity of amplification. The primer sequences used for qPCR are as follows: IL1β (5’-CCACCTCAATGGACAGAATATCA-3’, 5’-CCCAAGGCCACAGGTATTT-3’), IL6 (5’-CCAGAGTCCTTCAGAGAGATACA-3’, 5’-CCTTCTGTGACTCCAGCTTATC-3’, TNFα (5’-TTGCTCTGTGAAGGGAATGG-3’, 5’-GGCTCTGAGGAGTAGACAATAAAG-3’). The reactions were done in quadruplicate. The mRNA level in the reaction mixture was normalized by a reference PCR with endogenous GAPDH, and gene expression was determined with the comparative CT method (2^−ΔΔ^CT method).

### 2.10 Protein isolation

At the end of incubation, the total cellular protein and nuclear extracts were prepared with a commercial Nuclear and Cytoplasmic Protein Extraction Kit (Beyotime Biotechnology CO., Shanghai, China). And the protein concentration of the cellular lysate was determined using a Bicinchoninic Acid (BCA) Protein Assay Kit (Dalian Meilun Biotechnology Co., Dalian, China) by absorption at 562nm.

### 2.11 Immunoblotting analysis

The cellular lysates were mixed with a 6 × loading buffer (Beyotime Institute of Biotechnology, Haimen, China) and heated at 95 °C for 5min. Twenty micrograms of protein were separated by SDS-PAGE and transferred to polyvinylidene fluoride membranes (Immun-Blot, Bio-RAD, USA). After being blocked with 5% nonfat dry milk in Phosphate-buffered saline (PBS) plus 0.1% Tween 20 (PBST), membranes were then incubated with primary antibody and horseradish peroxidase-conjugated secondary antibodies, respectively. Signal detection was performed using enhanced chemiluminescence with the G: BOX Chemi XRQ (Syngene, USA) detection system. Analysis of band intensity was performed using ImageJ software. Antibodies used for immunoblot analysis are listed in Table 1.

**Table 1.** Antibodies used for experiments.

| Antibody | Vendor | Catalog no. | Working dilution |
| --- | --- | --- | --- |
| Mouse anti-GAPDH | ProteinTech | 60004-1-Ig | 1:50000 |
| Mouse anti-Lamin B1 | Beyotime Bio. Ld. | AF1408 | 1:1000 |
| Rabbit anti-CAT | ProteinTech | 21260-1-AP | 1:10000 |
| Rabbit anti-PPARγ | ProteinTech | 16643-1-AP | 1:5000 |
| Rabbit anti-PTGS2 | ProteinTech | 12375-1-AP | 1:2000 |
| Rabbit anti-IκB | ProteinTech | 10268-1-AP | 1:2500 |
| Rabbit anti-phospho-p65 | Beyotime Bio. Ld. | AF5881 | 1:1000 |

### 2.12 Statistical Analysis

Statistical differences between the control group and treatment groups were determined using an independent, two-tailed Student’s t-test. Experimental data are presented as mean±SD (standard deviation). Differences were considered significant when P was less than 0.05. All experiments were repeated three times independently.

## 3. Results

### 3.1 Active Ingredients in QP and related Targets

To reduce the extreme complexity of both chemical components and action mechanisms, the active ingredients of QP were screened by a combined pharmaceutical parameter: oral bioavailability (OB) >30% and drug-likeness (DL) >0.12. A total of 18 active ingredients of QP were selected, including 15 ingredients from Chuanqiong and 3 ingredients from Danggui (Table 1). And a total of 95 related targets were retrieved from the TCMSP database, and were subject to further analysis (Figure 1C).

**Figure 1.**
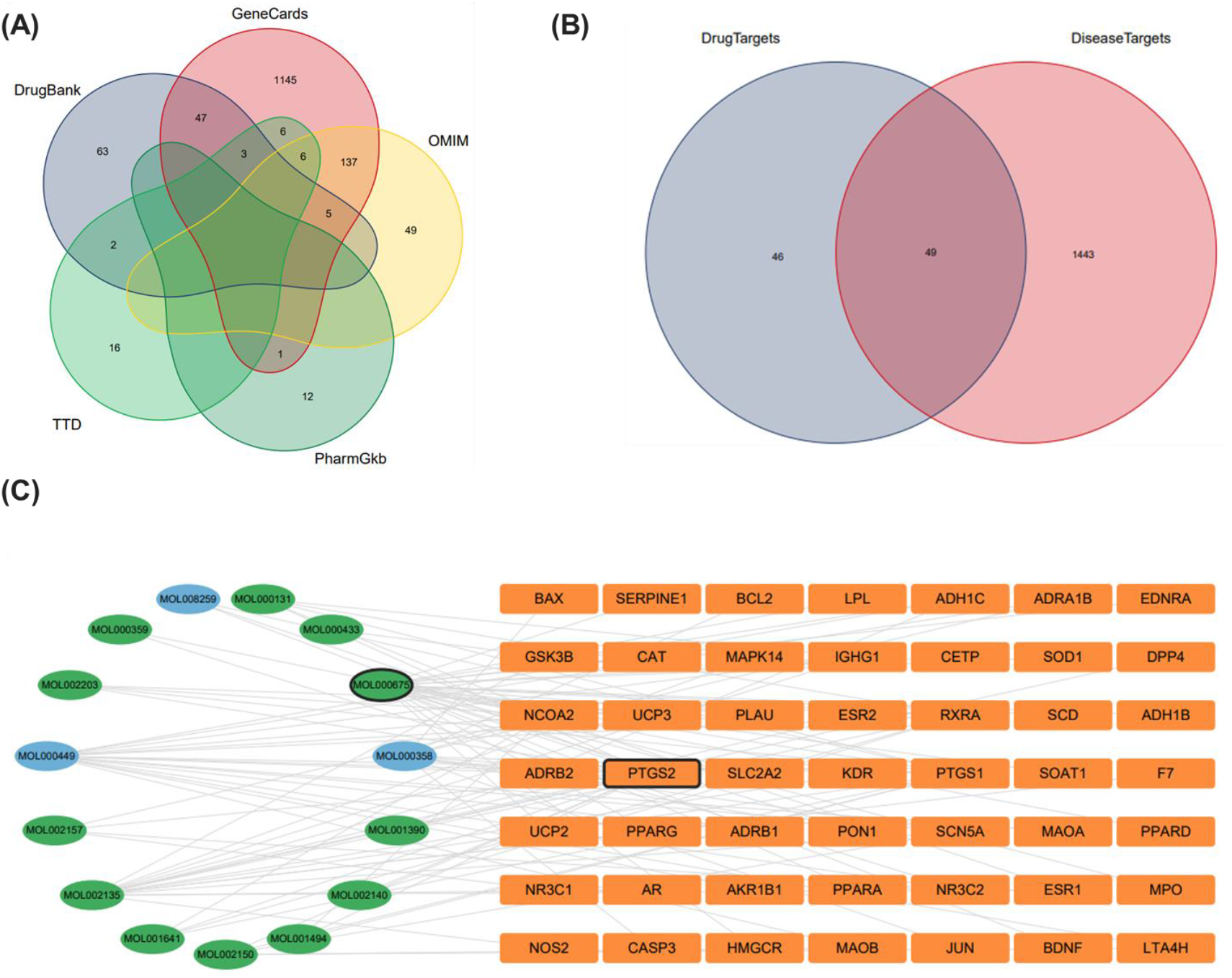
Construction of ingredient-target-disease network. (A) The Venn diagram of AS targets. (B) Venn plot showing the intersections of QP and AS-related genes. (C) The network of Ingredient-target-disease. the ring on the left represented the Ingredient and the rectangle on the right represented intersected genes.

### 3.2 Prediction of potential targets for AS

The potential targets for AS were established by retrieving related data from five public genetic and clinical databases with the keyword: AS. Thus, 1714 targets were obtained, including 1350 targets from GeneCards, 197 targets from OMIM, 13 targets from PharmGKB, 34 from TTD, and 120 targets from Drugbank. After screening out the duplicate data, a total of 1492 target genes are subject to further analysis (Figure 1A). Interaction analysis between QP targets and AS targets is shown in Figure 1B.

### 3.3 Ingredient-Target-Disease Network

To identify the pharmaceutical potentiality of QP against AS, a Venn diagram analysis was performed, which indicates the intersection between the target genes of QP and AS. A total of 49 target genes were identified. which are related to 15 active ingredients of QP. A network between the active ingredients and the intersected target genes was also constructed by Cytoscrape 3.8 software, and this graphical presentation elucidates the pharmaceutical potentiality of QP against AS (Figure 1C).

### 3.4 PPI Network Construction and Key Targets

To further understand the pharmaceutical potentiality of QP, the functional connectivity among these target proteins was analyzed by topological analyses. These target proteins were searched in the public STRING database, and a protein–protein interaction (PPI) network was constructed, wherein A total of 956 nodes and 1753 edges were defined after ruling out disconnected nodes (Figure 2A). To identify the hub proteins, the short tabular text was processed for two consecutive screenings with CytoNAC based on six topological factors: BC, CC, DC, EC, LAC, and NC. The values for screenings are listed in Table 3. Thus, a total of 15 hub proteins were identified, including CAT, NR3C1, PPARG, BDNF, SOD1, CASP3, PPARA, PTGS2, JUN, ESR1, NCOA2, RXRA, SERPINE1, NOS2, LPL (Figure 2B). And these proteins were considered for further verification by an LPS-induced inflammatory model *in vitro*.

**Figure 2.**
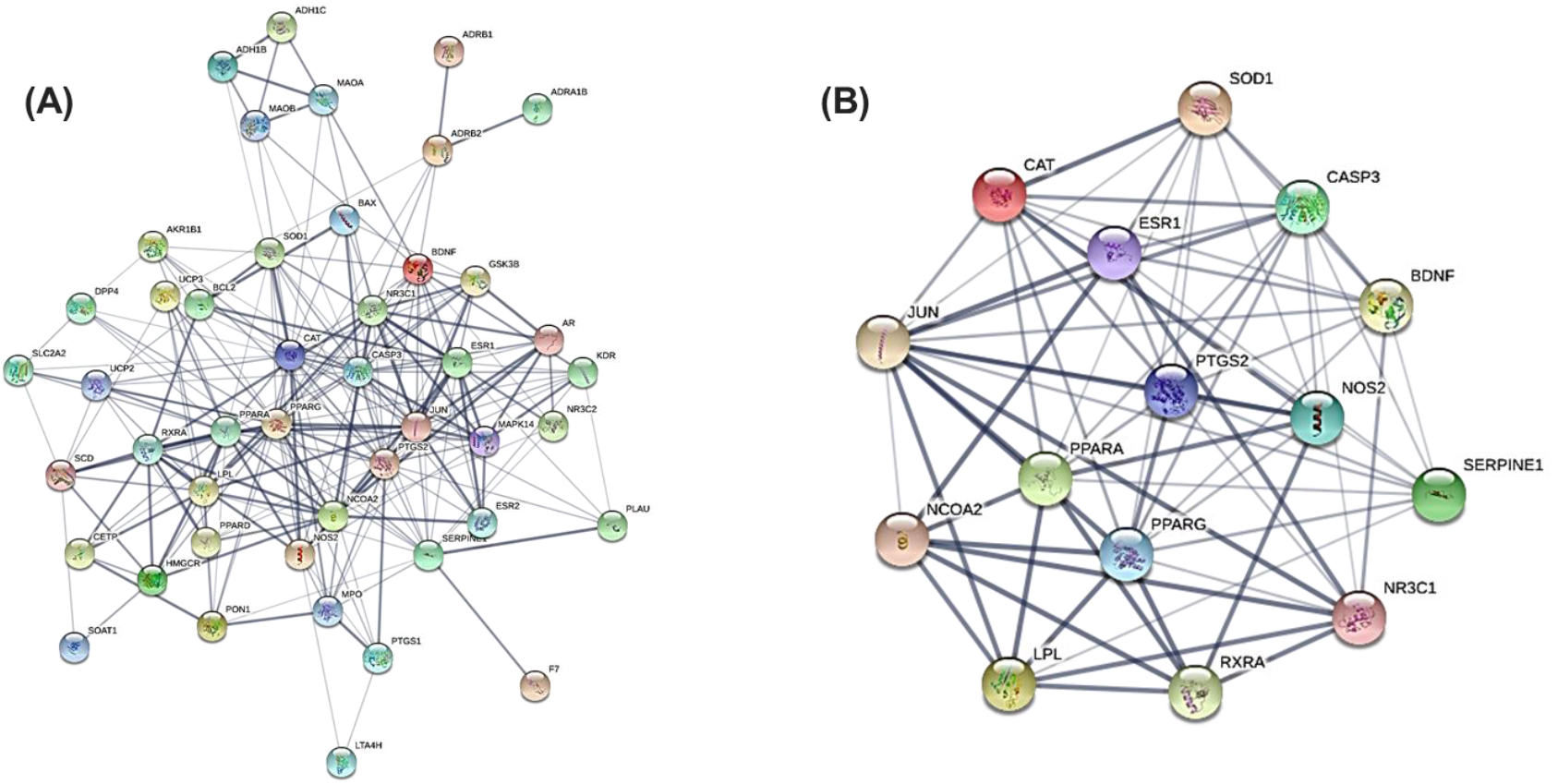
PPI network and cluster analysis of the intersected protein targets. (A) PPI network analysis with the intersected protein targets between QP and AS. (B) Hub network of protein targets, which was identified from (A) based on BC, CC, DC, EC, LAC, NC. BC.

**Table 2.** Active ingredients from QP.

| Herb Name | Mol ID | Molecule Name | OB (%) | DL |
| --- | --- | --- | --- | --- |
| *Ligusticum chuanxiong hort* | MOL001390 | 49070_FLUKA | 85.51 | 0.12 |
|  | MOL002153 | 1H-Cycloprop(e)azulen-7-ol, decahydro-1,1,7-trimethyl-4-methylene-, (1aR-(1aalpha,4aalpha,7beta,7abeta,7balpha))- | 82.33 | 0.12 |
|  | MOL000433 | FA | 68.96 | 0.71 |
|  | MOL002150 | 1-Acetyl-beta-carboline | 67.12 | 0.13 |
|  | MOL002140 | Perlolyrine | 65.95 | 0.27 |
|  | MOL002186 | Aromadendrene oxide 2 | 65.10 | 0.14 |
|  | MOL002151 | senkyunone | 47.66 | 0.24 |
|  | MOL002157 | wallichilide | 42.31 | 0.71 |
|  | MOL001494 | Mandenol | 42.00 | 0.19 |
|  | MOL001641 | METHYL LINOLEATE | 41.93 | 0.17 |
|  | MOL000131 | EIC | 41.90 | 0.14 |
|  | MOL002135 | Myricanone | 40.60 | 0.51 |
|  | MOL000359 | sitosterol | 36.91 | 0.75 |
|  | MOL000675 | oleic acid | 33.13 | 0.14 |
|  | MOL002203 | Exceparl M-OL | 31.90 | 0.16 |
| *Angelica sinensis* | MOL000449 | Stigmasterol | 43.83 | 0.76 |
|  | MOL000358 | beta-sitosterol | 36.91 | 0.75 |
|  | MOL008259 | 2,6-di(phenyl)thiopyran-4-thione | 69.13 | 0.15 |

**Table 3.** The topological parameters of hub targets.

| Name | Betweenness | Closeness | Degree | Eigenvector | LAC | Network |
| --- | --- | --- | --- | --- | --- | --- |
| PPARG | 349.4379 | 0.703125 | 54 | 0.291382 | 15.40741 | 37.24284 |
| CAT | 251.9959 | 0.642857 | 46 | 0.264028 | 15.65217 | 30.48418 |
| PTGS2 | 141.8851 | 0.642857 | 46 | 0.284782 | 18.26087 | 33.99031 |
| CASP3 | 105.3921 | 0.625 | 44 | 0.279289 | 18.36364 | 32.51118 |
| JUN | 93.73429 | 0.625 | 44 | 0.284919 | 19.09091 | 32.72603 |
| PPARA | 133.6271 | 0.625 | 42 | 0.231545 | 12.7619 | 24.62783 |
| BDNF | 124.5631 | 0.6 | 34 | 0.219684 | 15.52941 | 21.78275 |
| ESR1 | 42.02752 | 0.555556 | 32 | 0.211942 | 15.75 | 21.93898 |
| SERPINE1 | 142.4751 | 0.56962 | 32 | 0.203175 | 12.75 | 17.79333 |
| NR3C1 | 97.17427 | 0.576923 | 30 | 0.189594 | 12.8 | 16.58962 |
| LPL | 43.99028 | 0.54878 | 28 | 0.146451 | 12 | 17.69751 |
| SOD1 | 18.4446 | 0.542169 | 24 | 0.181194 | 13.66667 | 15.71635 |
| NCOA2 | 48.23114 | 0.529412 | 24 | 0.138128 | 9.333333 | 11.66501 |
| NOS2 | 70.95393 | 0.535714 | 24 | 0.169464 | 12.66667 | 15.72684 |
| RXRA | 40.5053 | 0.523256 | 22 | 0.109717 | 8.727273 | 10.70634 |

### 3.5 Functional Enrichment Analysis

To investigate the functional activities of QP in signaling pathways or biological processes, the KEGG and GO enrichment analyses were performed based on the potential target proteins of QP. The KEGG analysis revealed that these proteins are involved in 90 signaling pathways (p<0.5), and the top 3 pathways are significantly related to neurodegeneration, chemical carcinogenesis, as well as lipid and atherosclerosis (Figure 3A). Moreover, GO enrichment analysis revealed that these proteins were tightly related to response to nutrient levels and extracellular stimulus, membrane raft, microdomain, and DNA-binding transcription factor binding (Figure 3B). These data show the potential activities of QP in anti-atherosclerosis.

**Figure 3.**
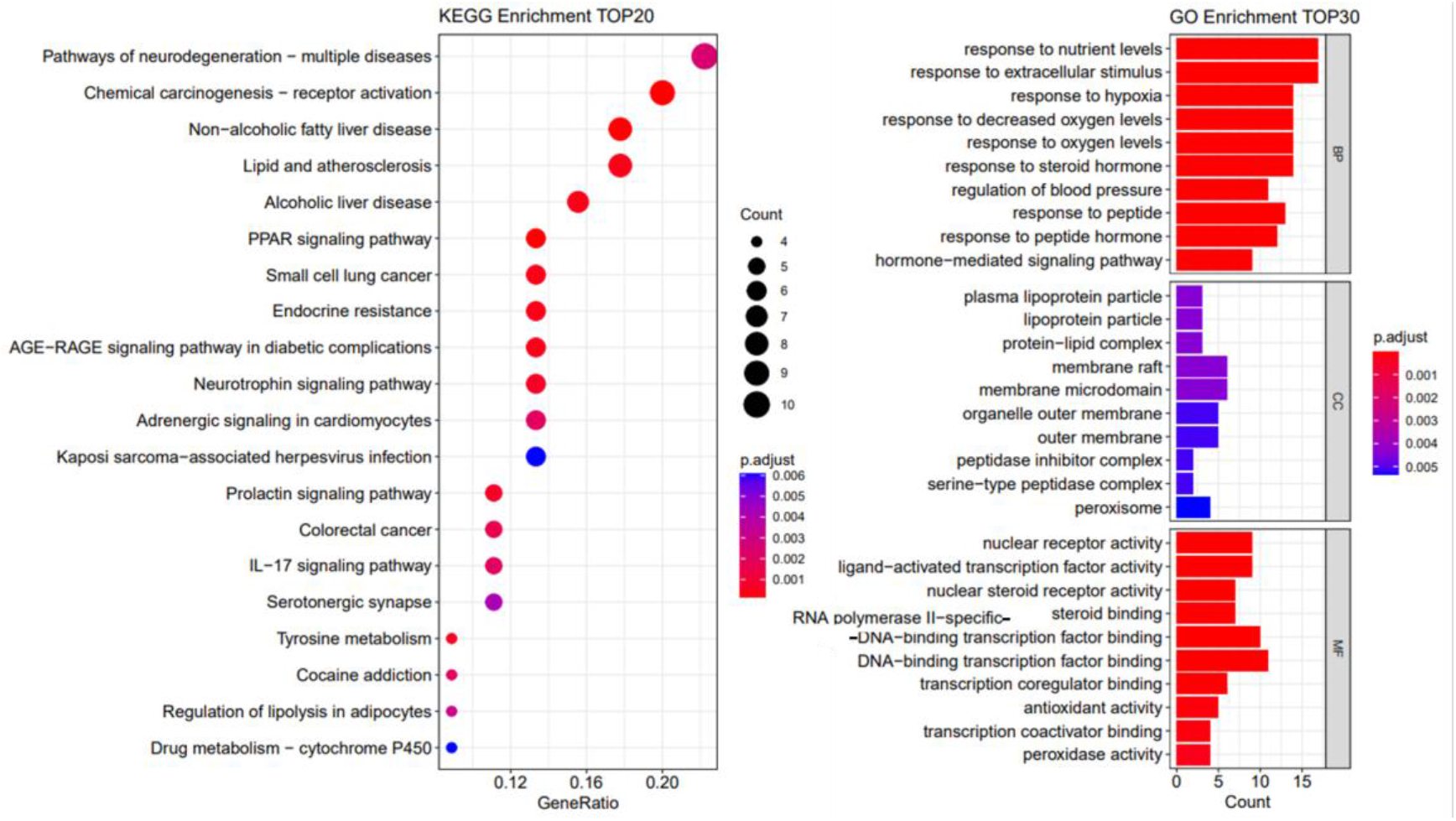
Gene Ontology (GO) and Kyoto Encyclopedia of Genes and Genomes (KEGG) functional analysis. (A) The top 30 enriched GO terms of intersected protein targets (B) The top 20 signaling pathways from KEGG analysis. BB: Biological process, CC: Cellular component, MF: Molecular function.

### 3.6 Effects Of QP Treatment On The Viability Of Macrophage Cells Line

To determine the QP effects on the viability of the macrophage, the RAW264.7 was incubated with different concentrations of QP for the indicated time, and the soluble formazan formation ability was assessed by WST-8 reduction assay (CCK-8 assay). Cells exposed to 0~4mg/ml QP exhibited a significant increase in WST-8 reduction after 6h treatment (p<0.05), whereas the increases, which were induced by treatment with 1mg/ml QP, were normalized generally after 12h treatment (Figure 4B). Moreover, QP treatment changed the morphology of RAW264.7, which was characterized by the increased number of vacuoles in the cytoplasm, in a concentration-dependent manner at 12h (Figure 4C). The vacuole formation might be reflective of altered biological processes in RAW264.7. Given that treatment with a higher concentration (>1mg/ml) of QP induced both deceased formazan formation and more visible vacuole, 0~1mg/ml of QP were chosen for further biochemical analysis. For more details regarding the characterization of formazan formation ability and morphology in RAW264.7 exposed to QP, see the supplementary Figure S1 and Figure S2.

**Figure 4.**
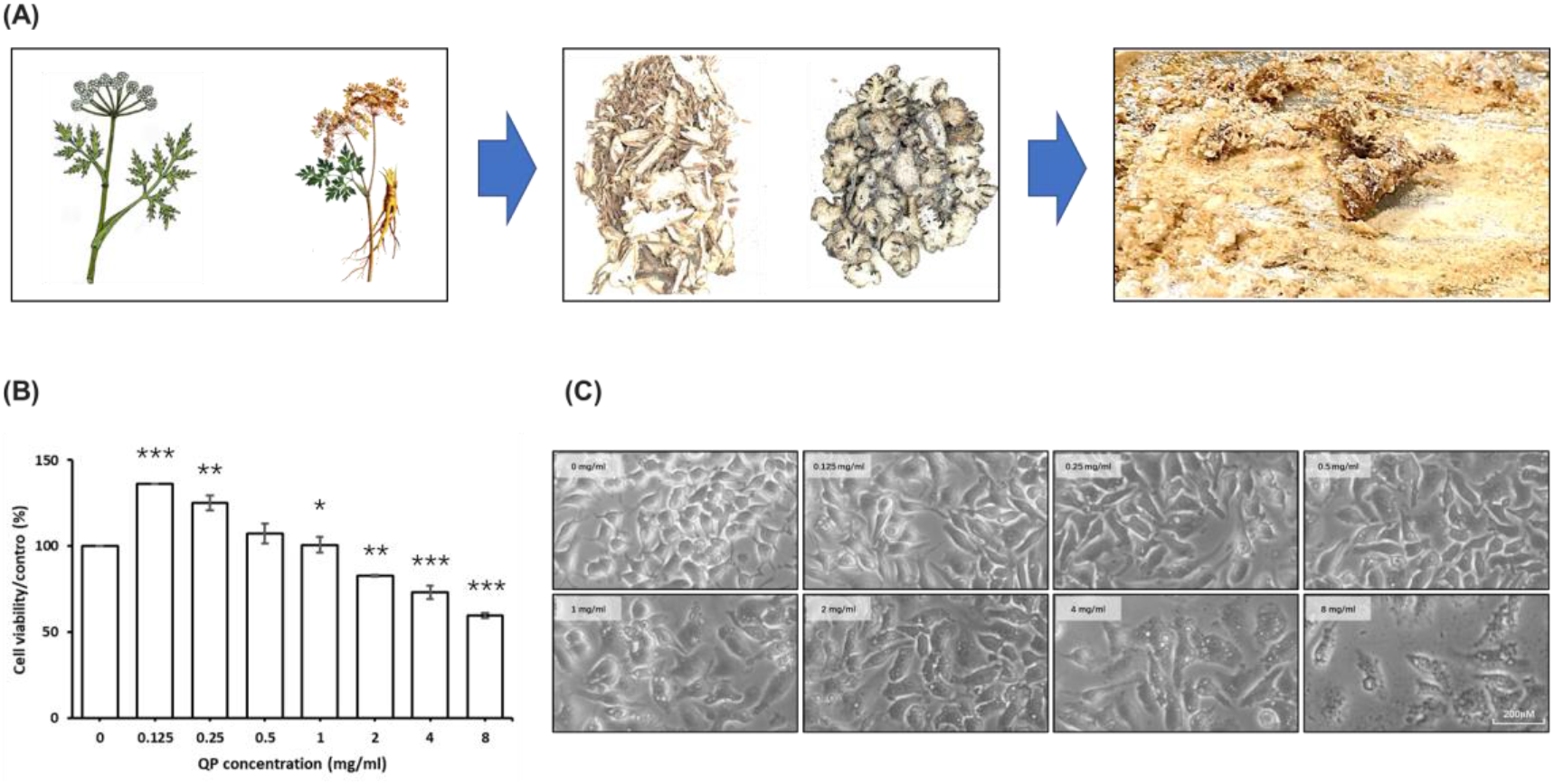
The effects of QP treatment on the biological activities of macrophage cells line. (A) Preparation of freeze-dried powders from water extraction of QP. The effects of QP treatment on the viability (B) and morphology (C) of RAW264.7.

**Figure 5.**
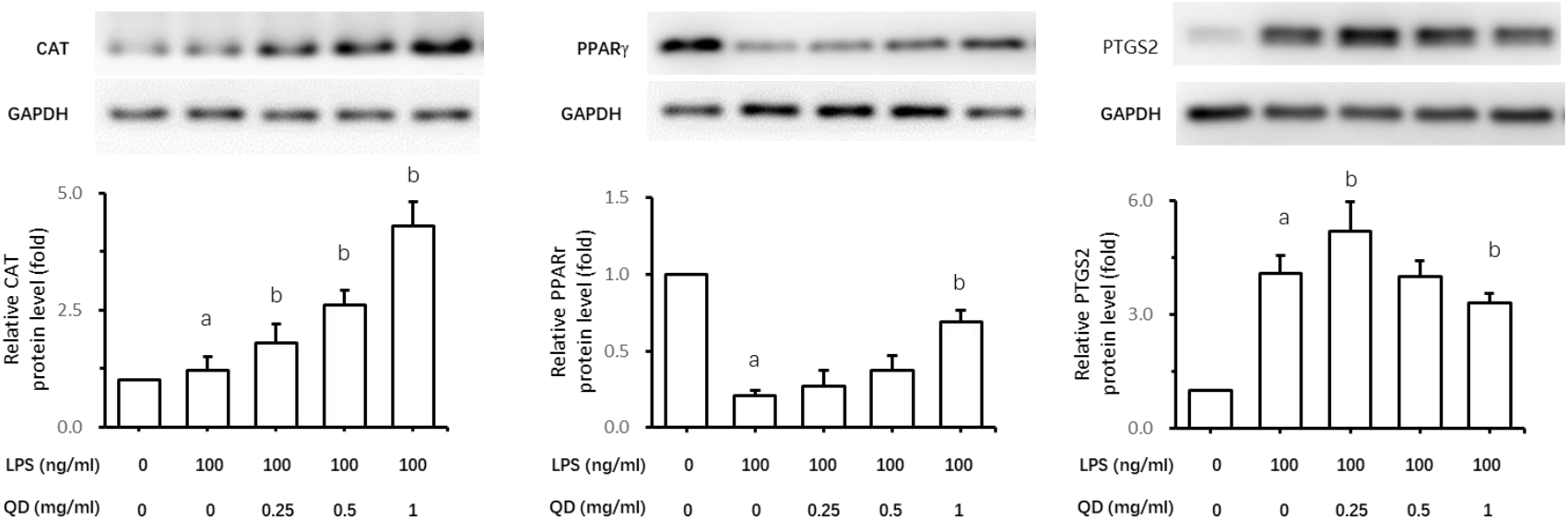
QP treatment affects the expression of CAT, PTGS2, and PPARγ. The RAW264.7 cells were treated with LPS (100ng/ml) in the presence or absence of QP for 12h, and the concentrations of protein level for CAT, PPAR4, and PTGS2 were detected by Western-blot assay. (a: p<0.05 compared to the control group; b: p<0.05 compared to the LPS group)

### 3.7 Effects Of QP On Peroxisome Proliferator-Activated Receptors Gamma (PPARγ), Catalase (CAT), Prostaglandin-Endoperoxide Synthase 2 (PTGS2)

To verify the pharmacological mechanism of QP against AS as predicted by the analysis *in silico*, the effects of QP on the expression of top 3 hub targets were measured using an inflammatory model in vitro, wherein LPS-stimulated macrophage display markers typical of AS-associated macrophage during AS pathogenesis. Treatment with QP increased the expression of CAT within LPS-treated RAW264.7 in a concentration-dependent manner. Treatment with QP (1mg/ml) also attenuated the increased expression of PTGS2, which is induced by LPS stimulus. And treatment with QP (1mg/ml) upregulated significantly the expression of PPARγ, but this effect was not observed in low-concentration groups. These data showed a general consistence with the prediction *in silico*.

### 3.8 Effects Of QP On The Inflammatory Mediates Secreted By Macrophage

Inflammation plays a central role in the pathogenesis of AS. And LPS-stimulated RAW264.7 is a well-established *in vitro* model of inflammation. We, therefore, investigate the effects of QP on the expression of inflammatory mediates in this model. Interestingly, QP exhibited an activity of anti-inflammation, evidenced by the reduced expression of inflammatory mediates including IL1, IL6, and TNFα, which are known markers for LPS-induced inflammation in RAW264.7 (Figure 6A). Moreover, this effect was accompanied by a suppression of NF-κB signaling pathways, because QP treatment alleviated the abnormal distribution of phosphorylated-p65 in the cytosol and nuclear, and increased the amount of IκB.

**Figure 6.**
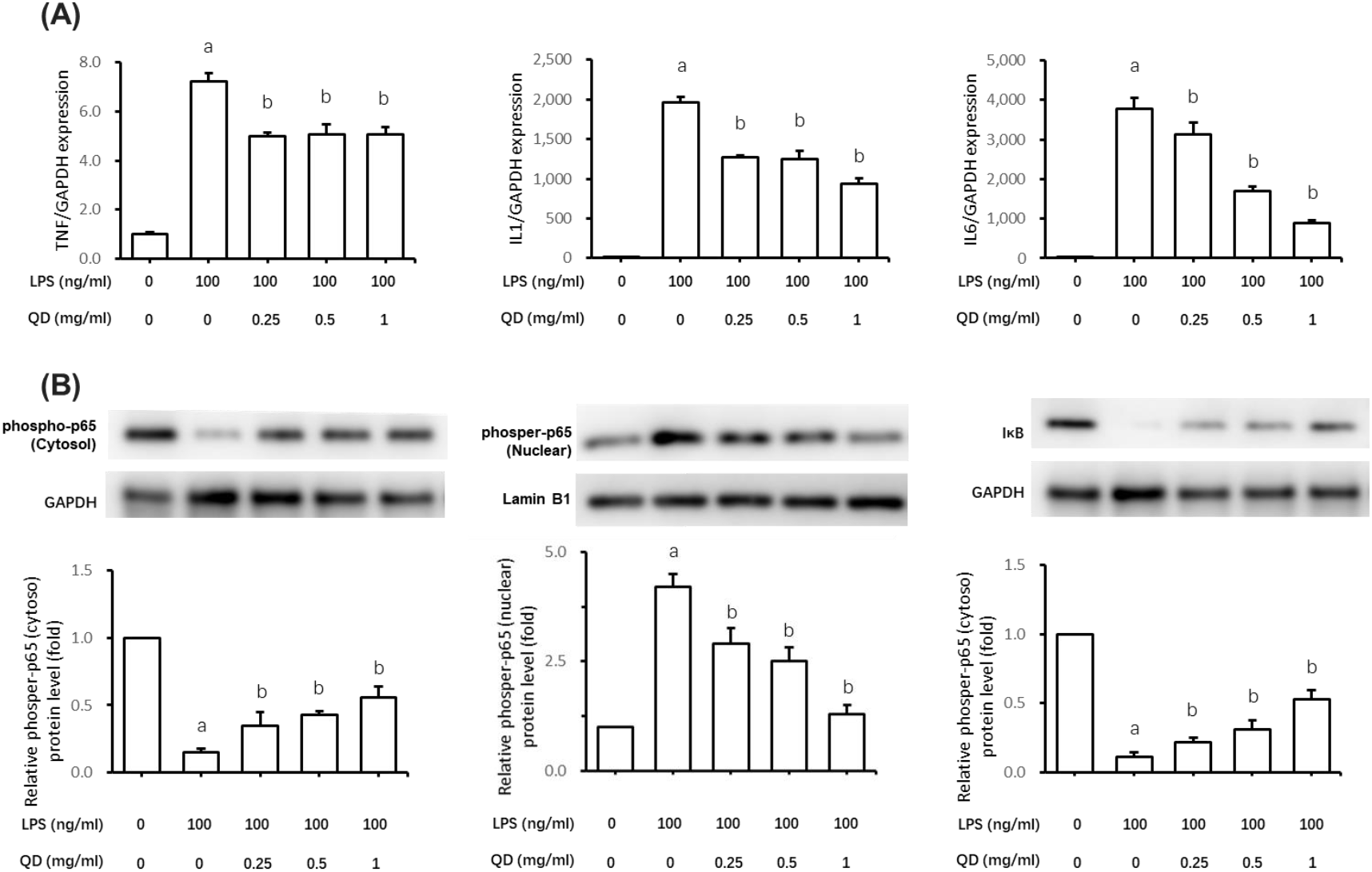
QP treatment exhibited anti-inflammatory effects on LPS-stimulated macrophage. (A) QP treatment induced a decrease in mRNA levels of TNFα, IL6, and IL1, compared with the LPS-stimulated group. (B) QP treatment suppressed NF-κb signaling activation induced by LPS stimulus. (a: p<0.05 compared to the control group; b: p<0.05 compared to the LPS group)

## 4. Discuss

Here, we have described the Network pharmacology analysis and functional verification of a Chinese herbal complex Qionggui Power (QP), which is composed of two herbs, Chuanqiong (*Ligusticum chuanxiong hort*) and Danggui (*Angelica sinensis*). The *in-silicon* analysis revealed active ingredients and hub molecular targets. These targets are highly enriched in the signaling pathway and biological processes highly related to AS pathogenesis. These findings are functionally significant because QP treatment exhibited general effects on the expression of core molecular targets, which are abnormally expressed in a macrophage model of inflammation. Moreover, *in vitro* experiments also showed an anti-inflammatory activity of QP, evidenced by the reduced expression of inflammatory mediators. Our data indicate a potential anti-atherosclerosis character of QP, which might contribute to further pharmaceutical development.

Screening of active ingredients from herbal medicine plays an important role in network pharmacology analysis. Among the most essential pharmacokinetic parameters for the screening of active ingredients are oral bioavailability ^[15]^ and drug-likeness ^[16]^. Oral bioavailability indicates the efficiency of ingredients reaching systemic blood circulation after oral administration, whereas drug-likeness suggests the resemblance of a particular molecule to existing drugs. By assessing quantitively physicochemical properties’ impact on molecular behavior, drug-likeness has been widely used to filter out undesirable compounds in the early phases of drug discovery ^[17, 18]^. The suggested drug screening criteria for TCMSP are OB ⩾ 20% and DL ⩾ 0.1. In the present study, the criteria were set as OB ⩾ 30% and DL ⩾ 0.12. The increased ratio might contribute to improving the screening of therapeutic bioactive molecules with more optimized pharmacokinetic and pharmaceutical properties.

Atherosclerosis involves pathophysiological alternations in medium to large arteries, which is largely driven by lipid abnormality and dysregulated inflammation ^[19]^. Treatments targeting lipid or inflammatory homeostasis ^[20]^ are useful for alleviating morbidity and mortality of atherosclerotic cardiovascular diseases. Our *in silico* data revealed 15 core pharmaceutical targets of QP, and these targets showed enrichment in multiple pathways, including Lipid and Atherosclerosis. The top 3 ranked proteins in the hub targets list are PPARγ, CAT, and PTGS2. PPARγ, along with PPARα and PPARβ, form a subfamily of the nuclear hormone receptor gene family ^[21]^. Previous studies have suggested that PPARα and PPARγ activation decreases AS progression not only by correcting metabolic disorders but also through direct effects on the vascular wall ^[22]^. The mechanism underlying this cardiovascular benefit involves lipid metabolism and inflammatory response. PTGS2 (also named COX-2), which is induced by cytokines and mitogens, contributes to inflammation, pain, angiogenesis, and cancer ^[23]^. And inhibition of PTGS2 results in reduced prostacyclin within the vessel wall, which exhibits a protective block on vascular inflammation and atherosclerosis ^[24]^. CAT can catalyze the decomposition of hydrogen peroxide into water and oxygen. The enhanced activity of CAT was revealed in foam cells obtained from atherosclerotic lesions from rabbit aortas. And a combined overexpression of CAT and SOD1 is linked to the alleviation of atherosclerosis ^[25]^. In the present study, QP treatment exhibited an atheroprotective effect, evidenced by an increase in PPARγ and CAT protein levels, as well as reduced PTGS2 expression. These data also correlate with the analysis by network pharmacology.

Macrophage is one of the key cell types that contributes to arterial inflammation. As the most abundant leukocyte in atherosclerotic lesions, macrophage decisively determines the inflammatory environment in the plague particularly by secreting inflammation-related cytokines, such as IL1, IL6, and TNFα ^[26, 27]^. Clinical agents that antagonize these cytokines have proven considerable effectiveness in reducing cardiovascular events ^[28, 29]^. Interestingly, macrophages exposed to QP exhibited a suppressed expression of IL1, IL6, and TNFα, which were induced by LPS, the most potent microorganism-derived inflammatory toxins. This suppression was accompanied by changes in the activities of the NF-κB signaling pathway, which is known as the critical regulator for inflammatory processes [30]. Such an anti-inflammatory effect was not reflected in the network pharmacology analysis of QP. Possible explanations include the paradigm of active ingredient screening from TCMSP. The higher screening criteria (OB⩾30% and DL⩾0.12.) facilitate the inclusion of desirable candidate components from the drug ingredients database for *in silico* analysis. However, this paradigm also increases the possibility of excluding active ingredients’ potential for anti-inflammation effects. Thus, an *in vivo* model of Atherosclerosis will be required for a global assessment of cardioprotective effects conferred by QP. Of note, this process may also complicate the validation of network pharmacology analysis due to the various cell types present in atherosclerotic lesions. The differential functions of target genes in these cells can be another consideration.

Collectively, our data provide several insights into the potency of QP in pharmaceutical development against atherosclerosis-related disease. The *in silico* analysis reveal active ingredients with favorable pharmacokinetic parameters (OB and DL) and the hub targets which are tightly involved in atherosclerosis pathogenesis and clinical intervents. Main predicted cardioprotective targets were validated in a macrophage model of inflammation, which comprises the major contributor to the formation of atherosclerotic lesions. Further work focusing on the cardioprotective effects *in vivo* is required to fully determine its value in pharmaceutics development.

## Author Contributions

Conceptualization, Y.W.; methodology, X.J., G.Z. Q.Z. and D.S.; software, Y.M.; validation, S.L.; formal analysis, X.J. and Y.M.; writing—original draft preparation, Y.M.; writing—review and editing, Y.W.; visualization, X.J.; supervision, Y.W.; project administration, Y.W. and S.L.; funding acquisition, Y.W. All authors have read and agreed to the published version of the manuscript.

## Data Availability Statement

All data generated or analyzed during this study are included in this article. Further inquiries can be directed to the corresponding author.

## References

[1] Mc Namara K, Alzubaidi H, Jackson JK: Cardiovascular disease as a leading cause of death: how are pharmacists getting involved? Integr Pharm Res Pract 2019;8:1–11.

[2] Sukumar S, Brodsky M, Hussain S, Yanek L, Moliterno A, Brodsky R, Cataland SR, Chaturvedi S: Cardiovascular disease is a leading cause of mortality among TTP survivors in clinical remission. Blood Adv 2022;6:1264–1270.

[3] Amini M, Zayeri F, Salehi M: Trend analysis of cardiovascular disease mortality, incidence, and mortality-to-incidence ratio: results from global burden of disease study 2017. BMC Public Health 2021;21:401.

[4] Ferranti SD de, Steinberger J, Ameduri R, Baker A, Gooding H, Kelly AS, Mietus-Snyder M, Mitsnefes MM, Peterson AL, St-Pierre J, Urbina EM, Zachariah JP, Zaidi AN: Cardiovascular Risk Reduction in High-Risk Pediatric Patients: A Scientific Statement From the American Heart Association. Circulation 2019;139:e603–e634.

[5] Bahekar AA, Singh S, Saha S, Molnar J, Arora R: The prevalence and incidence of coronary heart disease is significantly increased in periodontitis: a meta-analysis. Am Heart J 2007;154:830–837.

[6] Assmann G, Schulte H, Eckardstein A von, Huang Y: High-density lipoprotein cholesterol as a predictor of coronary heart disease risk. The PROCAM experience and pathophysiological implications for reverse cholesterol transport. Atherosclerosis 1996;124:S11–S20.

[7] Flora GD, Nayak MK: A Brief Review of Cardiovascular Diseases, Associated Risk Factors and Current Treatment Regimes. Curr Pharm Des 2019;25:4063–4084.

[8] Bäck M, Yurdagul A, Tabas I, Öörni K, Kovanen PT: Inflammation and its resolution in atherosclerosis: mediators and therapeutic opportunities. Nat Rev Cardiol 2019;16:389–406.

[9] Wang G, Dai G, Song J, Zhu M, Liu Y, Hou X, Ke Z, Zhou Y, Qiu H, Wang F, Jiang N, Jia X, Feng L: Lactone Component From Ligusticum chuanxiong Alleviates Myocardial Ischemia Injury Through Inhibiting Autophagy. Front Pharmacol 2018;9:301.

[10] Zhang Y, Ma C, He L, Liao L, Guo C, Wang C, Gong L, Zhou H, Fu K, Peng C, Li Y: Tetramethylpyrazine Protects Endothelial Injury and Antithrombosis via Antioxidant and Antiapoptosis in HUVECs and Zebrafish. Oxid Med Cell Longev 2022;2022:2232365.

[11] Wang J, Wang L, Zhou H, Liang X-D, Zhang M-T, Tang Y-X, Wang J-H, Mao J-L: The isolation, structural features and biological activities of polysaccharide from Ligusticum chuanxiong: A review. Carbohydr Polym 2022;285:118971.

[12] Pulendran B, S Arunachalam P, O’Hagan DT: Emerging concepts in the science of vaccine adjuvants. Nat Rev Drug Discov 2021;20:454–475.

[13] Tang J, Aittokallio T: Network pharmacology strategies toward multi-target anticancer therapies: from computational models to experimental design principles. Curr Pharm Des 2014;20:23–36.

[14] Hopkins AL: Network pharmacology: the next paradigm in drug discovery. Nat Chem Biol 2008;4:682–690.

[15] Xu X, Zhang W, Huang C, Li Y, Yu H, Wang Y, Duan J, Ling Y: A novel chemometric method for the prediction of human oral bioavailability. Int J Mol Sci 2012;13:6964–6982.

[16] Yamanishi Y, Kotera M, Kanehisa M, Goto S: Drug-target interaction prediction from chemical, genomic and pharmacological data in an integrated framework. Bioinformatics 2010;26:i246–54.

[17] Hu Q, Feng M, Lai L, Pei J: Prediction of Drug-Likeness Using Deep Autoencoder Neural Networks. Front Genet 2018;9:585.

[18] Liu H, Wang J, Zhou W, Wang Y, Yang L: Systems approaches and polypharmacology for drug discovery from herbal medicines: an example using licorice. J Ethnopharmacol 2013;146:773–793.

[19] Malekmohammad K, Bezsonov EE, Rafieian-Kopaei M: Role of Lipid Accumulation and Inflammation in Atherosclerosis: Focus on Molecular and Cellular Mechanisms. Front Cardiovasc Med 2021;8:707529.

[20] Soehnlein O, Libby P: Targeting inflammation in atherosclerosis - from experimental insights to the clinic. Nat Rev Drug Discov 2021;20:589–610.

[21] Tyagi S, Gupta P, Saini AS, Kaushal C, Sharma S: The peroxisome proliferator-activated receptor: A family of nuclear receptors role in various diseases. J Adv Pharm Technol Res 2011;2:236–240.

[22] Duval C, Chinetti G, Trottein F, Fruchart J-C, Staels B: The role of PPARs in atherosclerosis. Trends Mol Med 2002;8:422–430.

[23] Kirkby NS, Lundberg MH, Wright WR, Warner TD, Paul-Clark MJ, Mitchell JA: COX-2 protects against atherosclerosis independently of local vascular prostacyclin: identification of COX-2 associated pathways implicate Rgl1 and lymphocyte networks. PLoS One 2014;9:e98165.

[24] Zaitone SA, Moustafa YM, Mosaad SM, El-Orabi NF: Effect of evening primrose oil and ω-3 polyunsaturated fatty acids on the cardiovascular risk of celecoxib in rats. J Cardiovasc Pharmacol 2011;58:72–79.

[25] Poznyak AV, Grechko AV, Orekhova VA, Chegodaev YS, Wu W-K, Orekhov AN: Oxidative Stress and Antioxidants in Atherosclerosis Development and Treatment. Biology (Basel) 2020;9.

[26] Moore KJ, Sheedy FJ, Fisher EA: Macrophages in atherosclerosis: a dynamic balance. Nat Rev Immunol 2013;13:709–721.

[27] Oikonomou E, Tsaplaris P, Anastasiou A, Xenou M, Lampsas S, Siasos G, Pantelidis P, Theofilis P, Tsatsaragkou A, Katsarou O, Sagris M, Vavuranakis M-A, Vavuranakis M, Tousoulis D: Interleukin-1 in Coronary Artery Disease. Curr Top Med Chem DOI: 10.2174/1568026623666221017144734.

[28] Barrett TJ: Macrophages in Atherosclerosis Regression. Arterioscler Thromb Vasc Biol 2020;40:20–33.

[29] Libby P: The changing landscape of atherosclerosis. Nature 2021;592:524–533.

[30] Lai CC, Nelsen B, Frias-Anaya E, Gallego-Gutierrez H, Orecchioni M, Herrera V, Ortiz E, Sun H, Mesarwi OA, Ley K, Gongol B, Lopez-Ramirez MA: Neuroinflammation Plays a Critical Role in Cerebral Cavernous Malformation Disease. Circ Res DOI: 10.1161/CIRCRESAHA.122.321129.

