## Supplemental Table 1 and Supplemental Figure 1 for "Integrating Network Pharmacology And Experimental Verification To Explore The Mechanism Of Qionggui Power Against Atherosclerosis"

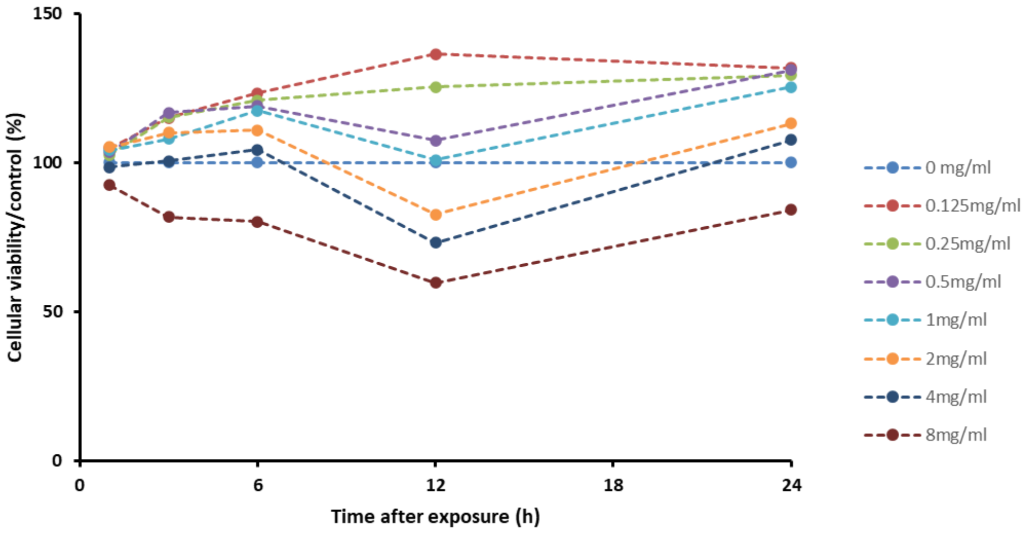


Figure S1: The viability of RAW264.7 after treatment with QP (0-8mg/ml) for indicated time (1h, 3h, 6h, 12h, and 24h).

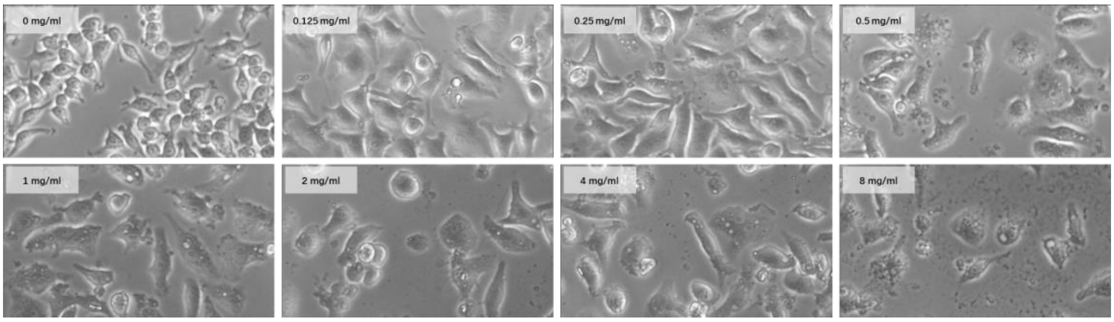

Figure S2: The morphology of RAW264.7 after treatment with QP for 24h
